# Ketamine attenuates habenula activity in response to aversive outcomes during Pavlovian learning

**DOI:** 10.64898/2026.02.08.704729

**Authors:** Erdem Pulcu, Sara Costi, Pilar Artiach-Hortelano, Chloe Wigg, Sorcha Hamilton, Marieke Martens, Rebecca P. Lawson, Rupert McShane, Philip Cowen, Susannah E. Murphy, Catherine J Harmer

**Affiliations:** Department of Psychiatry, University of Oxford, Oxford, OX3 7JX, UK; Oxford Health Foundation Trust, Warneford Hospital, Oxford, OX3 7JX, UK; Depression and Anxiety Center for Discovery and Treatment, Department of Psychiatry, Icahn School of Medicine at Mount Sinai, New York, NY, 10029 USA; Department of Psychology, University of Bath, Bath, UK; Department of Psychology, University of Cambridge, Cambridge, CB2 3EB

## Abstract

Ketamine is an NMDA receptor antagonist with rapid-antidepressant properties when administered at a sub-anesthetic dose. Preclinical models indicate that a direct injection of ketamine into lateral habenula (Hb), a small midbrain structure with an evolutionarily preserved role in aversive learning across mammals, can rapidly relieve depression-like behavior. However, there is limited evidence to explain how ketamine acts on the function of the human habenula. In a translational computational neuroscience study, 70 healthy adult volunteers were randomised in a 1:1 ratio to receive ketamine or placebo (NaCl 0.9%). We used an aversive Pavlovian conditioning paradigm combined with 7-Tesla functional neuroimaging to show that ketamine attenuates habenula response during aversive stimuli expectations and outcomes 24 hours post-infusion. We further present preliminary evidence suggesting that when aversive learning occurs after ketamine infusion, reduced habenula activity during the learning process may lead to downstream effects that diminish the aversive impact of negative affective memories. These findings provide translational support for preclinical models of ketamine’s mechanisms in humans.

## Introduction

Ketamine is an NMDA receptor antagonist with rapid antidepressant properties when administered at a sub-anaesthetic dosage^1-3^. Although the clinical efficacy of ketamine is reasonably well established^4,5^, the cognitive and neurobiological bases of its rapid antidepressant effects in the human brain are less characterised^6^.

Evidence from preclinical mouse models indicates that depression-like behaviours are associated with heightened burst firing of neurons in the lateral habenula^7^, a small brain region that is evolutionarily preserved across species^8,9^. This characteristic burst-firing signature in lateral habenula (Hb) neurons, along with the associated behavioural stress response, can be normalised by direct injection of ketamine into the region. Mechanistic insights from this preclinical model have described a key candidate neurobiological pathway that could underpin ketamine’s rapid antidepressant action in humans. Specifically, ketamine seems to act by suppressing burst firing of lateral habenula neurons, an action which is extended in time through the trapping of ketamine in NMDA receptor ion channels^10^. Here, we investigate the translational relevance of this rodent model, using an experimental medicine framework to determine how ketamine modulates activity in the human habenula.

In computational neuroscience, the role of the habenula has been documented across a number of influential studies conducted in monkeys^11,12^ and humans^13-15^, collectively suggesting that the habenula plays a central role in aversive learning^16-19^. In humans, a functional magnetic resonance imaging (fMRI) study measuring activity in the habenula during the anticipation of positive and negative outcomes (e.g. such as anticipation of electric pain delivery to the hand) demonstrated that activity in this region positively correlates with the expected values of aversive events during Pavlovian/observational learning^14^. These previous monkey neurophysiology and human fMRI studies clearly demonstrated that aversive learning paradigms provide experimental probes that engage habenula activity. In the present study, we used an aversive Pavlovian learning task (Figure 1) to probe the human habenula and to understand how ketamine modulates its activity during the learning process.

**Figure 1.**
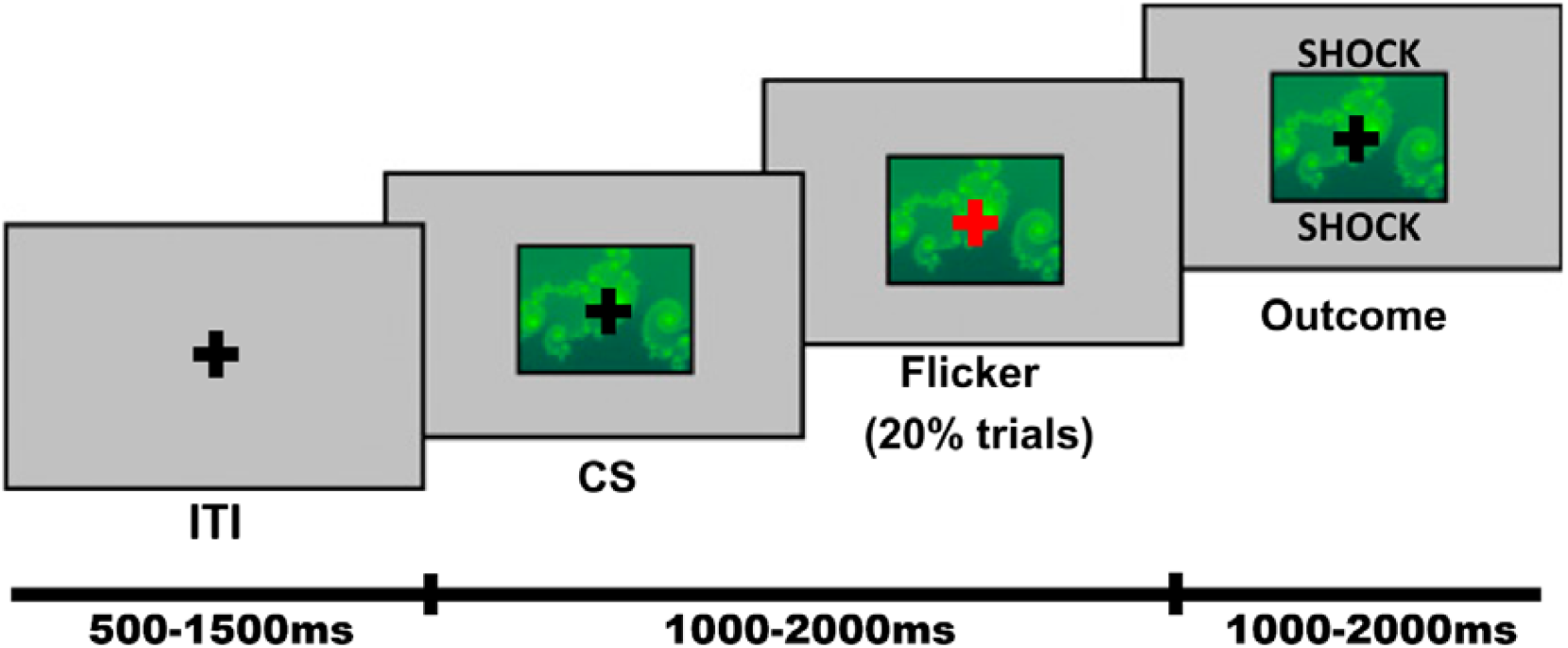
Timeline of the Pavlovian learning task. The participants were asked to form associations between abstract fractals and 4 different types of outcomes (winning or losing money, neutral, electric pain) through observational learning. The task consisted of 3 distinct blocks and each block contained 7 different fractals, i.e. 21 in total. Win, loss and shock fractals had high and low probability outcomes levels, i.e. 80% versus 20%; the neutral fractals were deterministically (i.e. 100% of the time) associated with null outcomes. In order to make sure that participants paid adequate attention to the observational learning task, 20% of the trials had red flickers superimposed on the fractals to which participants were asked to respond with a button press as quickly as possible. The mean duration of each epoch of the task is shown at the bottom the task timeline. Note that all task epochs were jittered, and durations were decorrelated to boost fMRI design efficiency. ITI: inter-trial interval (timing range 500-1500ms, uniformly distributed). CS: conditioned stimulus (timing range 1000-2000ms, uniformly distributed). Outcome duration 1000-2000ms, uniformly distributed. Figure adapted from Lawson et al., 2014 PNAS.

In monkey neurophysiology, it has been demonstrated that the habenula particularly tracks the learning signals associated with the most aversive stimuli-outcome pairs in the environment^12^. In humans, previous work also demonstrated a stronger response in the habenula to cues predicting primary punishments such as delivery of painful stimuli to the hand, relative to cues preceding secondary punishments such as losing money^14^. Here, we report findings from a cohort of 66 healthy volunteers without any previous psychiatric history, who were administered ketamine intravenously (IV) at a sub-anaesthetic dosage of 0.5mg per kilogram [of participant weight]^5^, and 24 hours later completed a Pavlovian aversive learning task while undergoing 7-Tesla functional neuroimaging. Pavlovian learning required participants to form associations between different types of conditioned stimuli (CS, i.e., visually distinct fractals as shown in Figure 1) and outcomes (win, loss, shock and null) through observation. We predicted that ketamine would attenuate habenula activity during the expectation phase as well as in response to peripheral delivery of electric pain (i.e. the outcome/pain delivery phase) during Pavlovian aversive learning, which formed our preregistered primary outcome measures for this study (clinicaltrials.gov identifier: NCT04850911). Our findings provide evidence that, 24h after administration, ketamine attenuates habenula activity in a similar fashion to that suggested by earlier preclinical models, and we further extend these findings by documenting potential downstream affective memory effects of reducing habenula activity following ketamine administration.

## Results

### fMRI results

This section characterises task-evoked activation at the whole-brain level and within the Hb region-of-interest (ROI) contralateral to shock stimulation, and then tests the effects of ketamine compared to placebo on these stimuli. All Hb ROI findings were conducted using a preregistered right Hb mask derived from the Pauli et al.^20^ atlas (probability ≥50%, binarized) and are reported as small volume correction (SVC), threshold-free cluster enhancement (TFCE), FWE-corrected within the ROI (*randomise*, N = 5000 permutations).

#### Pavlovian Learning evoked right Habenula Activity During Aversive Processing

Across all participants, activity during conditioned stimuli (CS) presentations in the dorsal cerebellum and medial temporal gyrus was positively correlated with the expected value of abstract fractals (**Figure 2A**), where these expected values were estimated for each individual through computational modelling (**see Materials and Methods**). In contrast, our primary focus was on regions exhibiting negative correlations with expected value signals *i.e*. those activated more to negatively perceived stimuli. These included the bilateral insula and a large, contiguous cluster extending posteroventrally from the thalamus to the habenula bilaterally (Fig. 2b).

**Figure 2.**
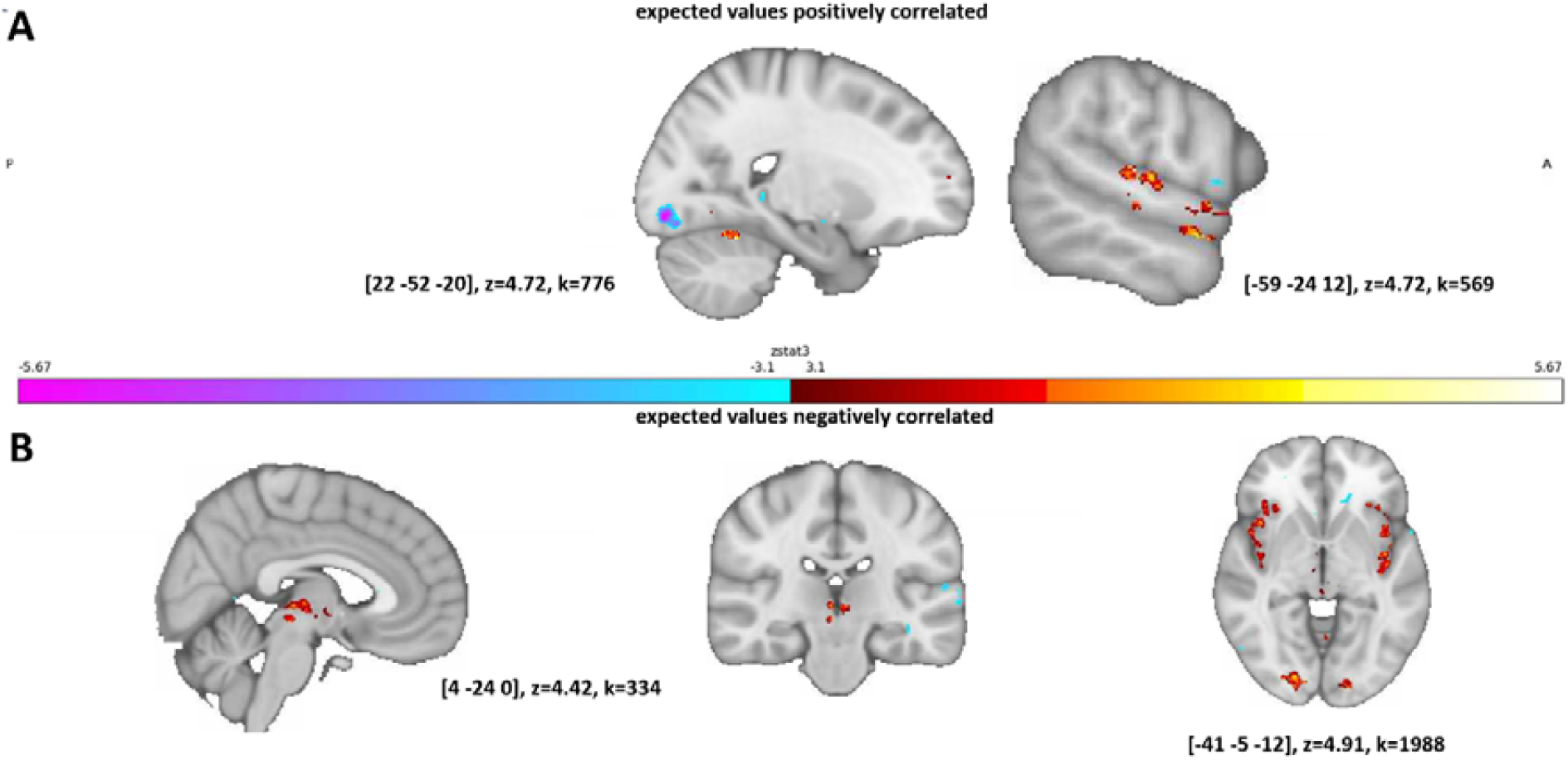
fMRI results, areas commonly activated across participants. (A) Expected values of the fractals were positively correlated with the activity in [both shown in sagittal view] dorsal aspect of the cerebellum (peak Montréal Neurological Institute (MNI) coordinates (XYZ): 22 -52 -20, z-value at peak: 4.72, cluster size (k): 776) and medial temporal gyrus (MNI: -59 24 12; z-max = 4.72; k = 569). (B) As expected, the task engaged regions implicated in aversive learning and pain processing such as insula [shown axially] (MNI: -41 -5 -12, z = 4.91, k = 1988) and a large cluster in midbrain [shown both sagittal and coronally] including the habenula bilaterally (MNI: 4 24 0, z = 4.42, k = 334). Colour bars represent z-values.

During US (i.e. outcome) presentations, there was robust whole-brain activation to the delivery of aversive (including shock) outcomes relative to baseline, as well as compared to positive stimuli, an effect that was also confirmed within the right Hb ROI (See Table S1 and S2 in the Supplementary Materials). Jointly, these regions define the whole-brain networks activated during CS and US presentations across all participants (irrespective of drug administration group).

#### Ketamine Attenuates Right Habenula Responses to the Expectation and Delivery of Aversive Stimuli

To investigate the effect of ketamine on right Hb response (i.e. contralateral hemisphere to the hand on which shocks were delivered) to expectations of aversive outcomes (**Figure 2B**), we conducted a permutation test via FSL’s *randomise* function within our preregistered a priori right Hb ROI. As hypothesised, the ketamine group showed significantly lower right Hb activity in response to aversive/negatively valued CS presentations on a positive to negative value spectrum, relative to placebo (MNI: [4 -24 4], p=0.023, Family-wise Error (FEW) corrected number of voxels:2, t-max value: 3.626), modulated by their perceived valence [based on a post-scan preference test results reported in Behavioural Results section below]. Bar plots of the parameter estimates within the right Hb ROI in each group are reported in **Figure 3A**. By contrast, no selective effects were observed when comparing each condition (shock, loss and win) individually (**Supplementary Figure 1**).

**Figure 3.**
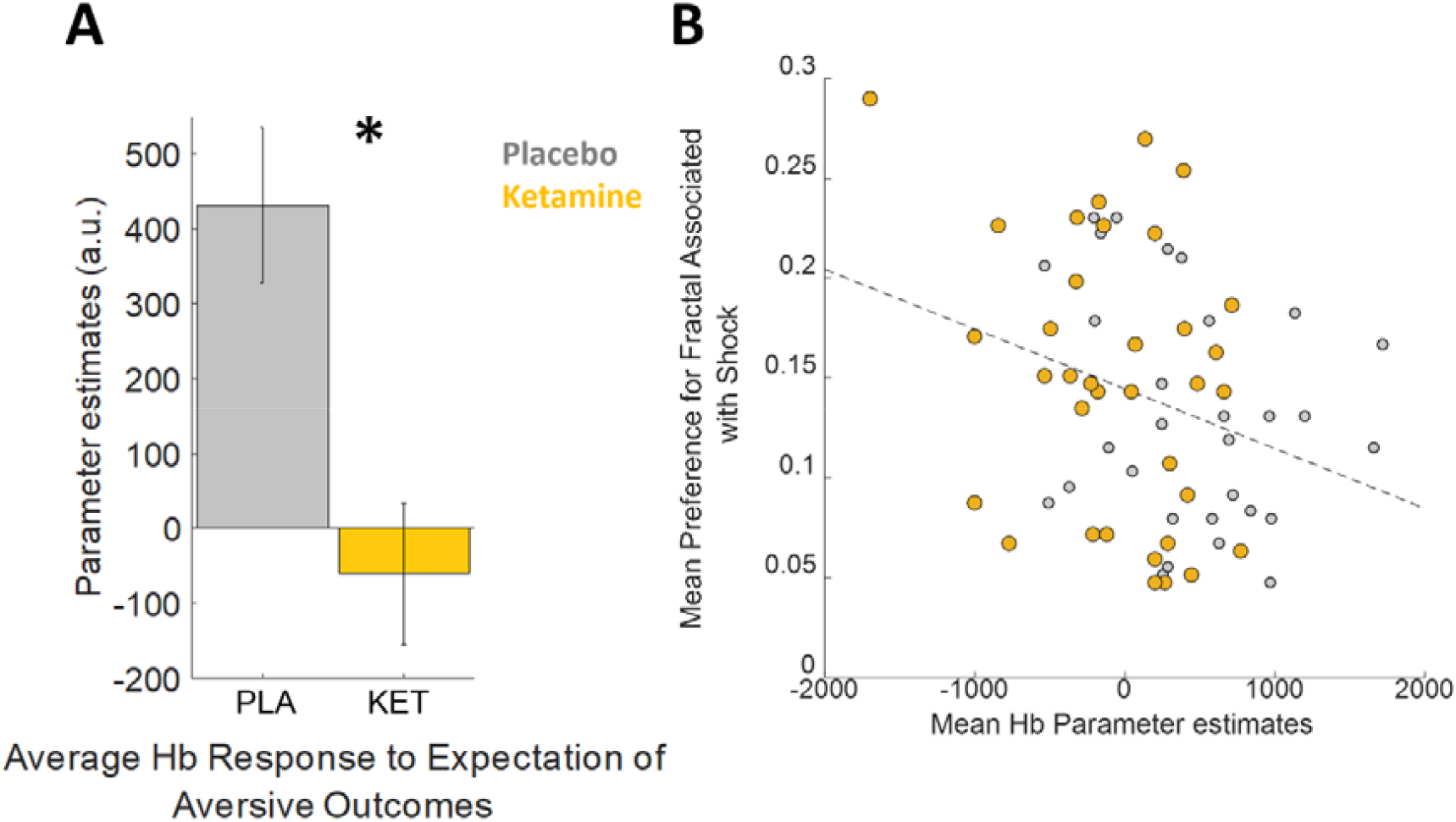
Habenula activity during the outcome expectation phase and subsequent choice preference for fractals associated with shock. (A) Subsequent between-group comparison confirms that ketamine attenuates habenula response to expectation of aversive outcomes (note that in the model-based analysis the same fMRI regressor linearly scales expectations of outcomes over an appetitive-aversive spectrum, *p<.05, FWE corrected) . (B) Average habenula activity correlates significantly negatively with the subsequent preference for fractals which were associated with electric pain outcomes (also the most aversive stimuli in the environment based on preference test results. Note that the preference test was administered approximately 30 minutes after the fMRI session and outside the scanner). Error bars designate ±1SEM.

Similarly, during the delivery of aversive stimuli, analysis of the right Hb ROI revealed a statistically significant cluster for the negative (shock + loss) > positive (win) contrast, with reduced activity in the ketamine group relative to placebo (22 voxels; MNI 4, -22, 2; p = 0.014, FWE-corrected; max t-stat = 2.25 **Figure 4A**). To further characterise this effect, we examined individual contrasts within the US. The loss > win contrast was greater in the placebo group relative to ketamine group (32 voxels; MNI: 4, –22, 2; p = 0.016, FWE-corrected; max t = 3.06; **Supplementary Figure 2**). Notably, the shock > win contrast was numerically higher in the placebo vs ketamine group, although this difference did not reach statistical significance (p = 0.089). At the whole-brain level, the delivery of aversive stimuli in the outcome phase did not elicit differential activation between ketamine and placebo. These results suggest that ketamine differentially modulates right Hb engagement during the processing of aversive versus positive outcomes in healthy volunteers, which is not specific to shock alone (**Figure 4B**).

**Figure 4.**
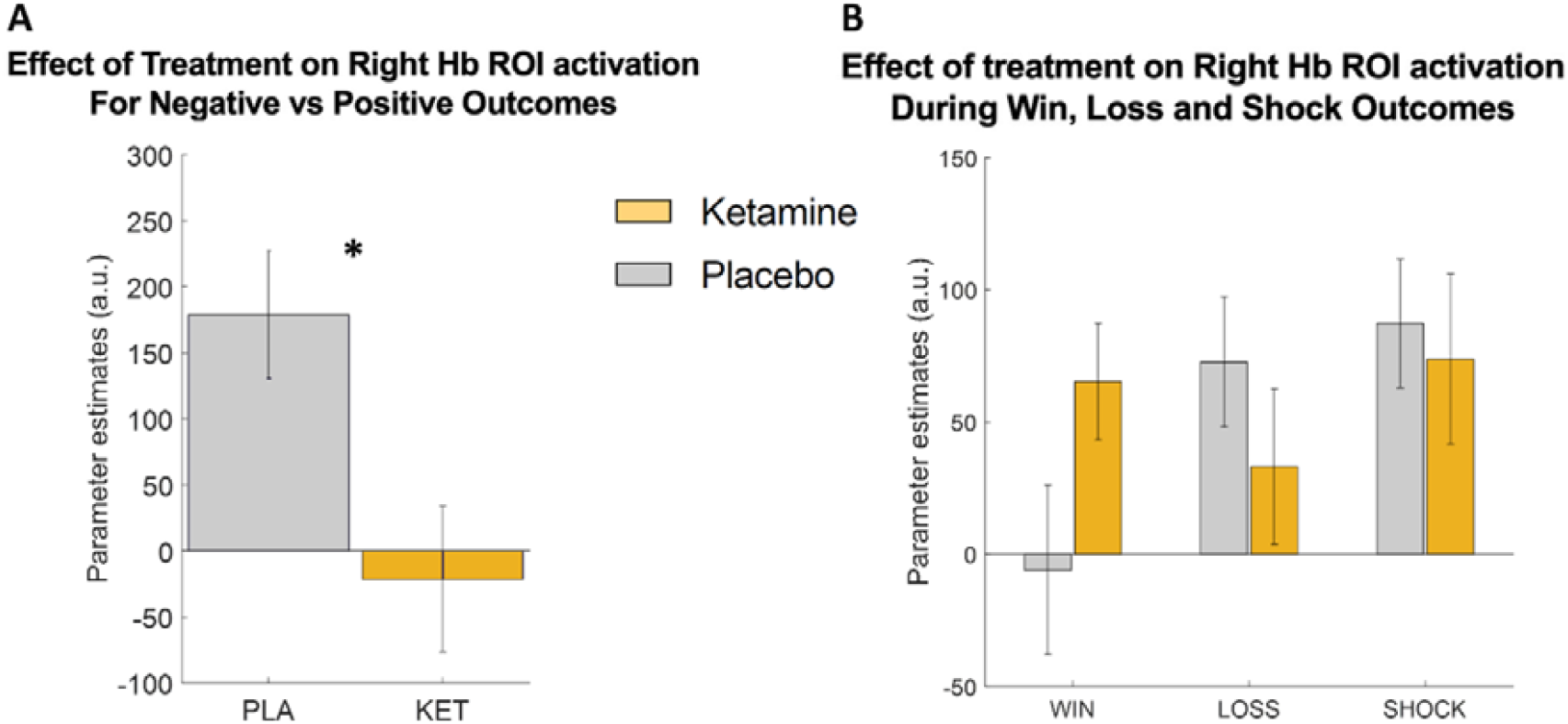
Ketamine attenuates right habenula response to delivery of aversive outcomes (US phase). (A) Ketamine group had significantly lower activity for the negative (i.e. shock+loss) versus positive (i.e. win) outcomes (*p=.014). Note that directionality of these effects remain stable in fMRI models with or without parametric modulators accounting for participants’ subjective preference between CS categories. (B) Right habenula (Hb) activity did not differ significantly between ketamine and placebo groups when examined separately for win (p=.067), loss (p=.31), or shock (p=.74) conditions [vs baseline]. Error bars designate ±1SEM.

### Behavioural results

Although the Pavlovian/observational learning component of our experimental design did not yield behavioural readouts directly indexing learning, there were two ancillary behavioural metrics: (i) in-scanner flicker detection response times (RTs; **Figure 1**), (ii) choice behaviour and RTs in a binary preference test between fractals previously observed during Pavlovian learning (i.e. a probe of memory-guided decision-making administered approximately 30 minutes later than the fMRI task and outside the scanner).

The drug administration groups were comparable in RTs for the in-scanner flicker-detection task and for RTs during the preference test. In the preference test, there was a main effect of valence on choice behaviour (F(3, 192)= 184.50, p<.001, **Supplementary Figure 3B**), with no main effect of drug administration group (F(1,64)=.066, p=.798), or valence x drug administration interaction (F(3,192)=.686, p=.562). Likewise, there was a main effect of valence on RTs during the preference test (F(3,192)= 154.13, p<.001), with slower responses to more negative CS associated stimuli (from win to shock, **Supplementary Figure 3C**), and no effect of treatment.

### Linking fMRI and behavioural results

First, we investigated whether right Hb activity during the CS presentation phase was related to behavioural responses (RTs) in the flicker-detection trials. Previous work^14^ suggested a significant negative association between Hb activity while anticipating shock outcomes and conditioned suppression (defined as slowing down of responses to flickers presented in shock vs neutral trials), and a significant positive relationship between Hb activity while expecting win outcomes and conditioned invigoration (defined as speeding up of responses to the flickers presented in win vs neutral trials). In the present cohort, however, neither of these relationships reached significance in the placebo group (n=31; all p>0.26).

Next, we investigated whether right Hb responses during the expectation phase of aversive outcomes (i.e. the CS presentation epoch on the task timeline) predicted subsequent choice behaviour in the binary preference test administered 30 minutes post-scanning. Both the CS/expectation phase (**Figure 3A**) and the binary preference test (**Supplementary Figure 3B**) engage the same cognitive process (estimating outcome probability), providing a natural link between these measures. Even after controlling for treatment group allocation (ketamine versus placebo) and subjective pain ratings (**Figure 5**), we observed a significant negative association between average right Hb activity to aversive CS presentations (**Figure 3A**) and subsequent preference for the shock-associated fractals in the preference test (r(62) =-0.279, p=0.025, shown in **Figure 3B**). Importantly, this negative relationship was observed in both ketamine (r=-0.287) and placebo (r=-0.208) groups, albeit nonsignificantly (both p>.0949), nevertheless ruling out a potential Simpson’s paradox^21^.

**Figure 5.**
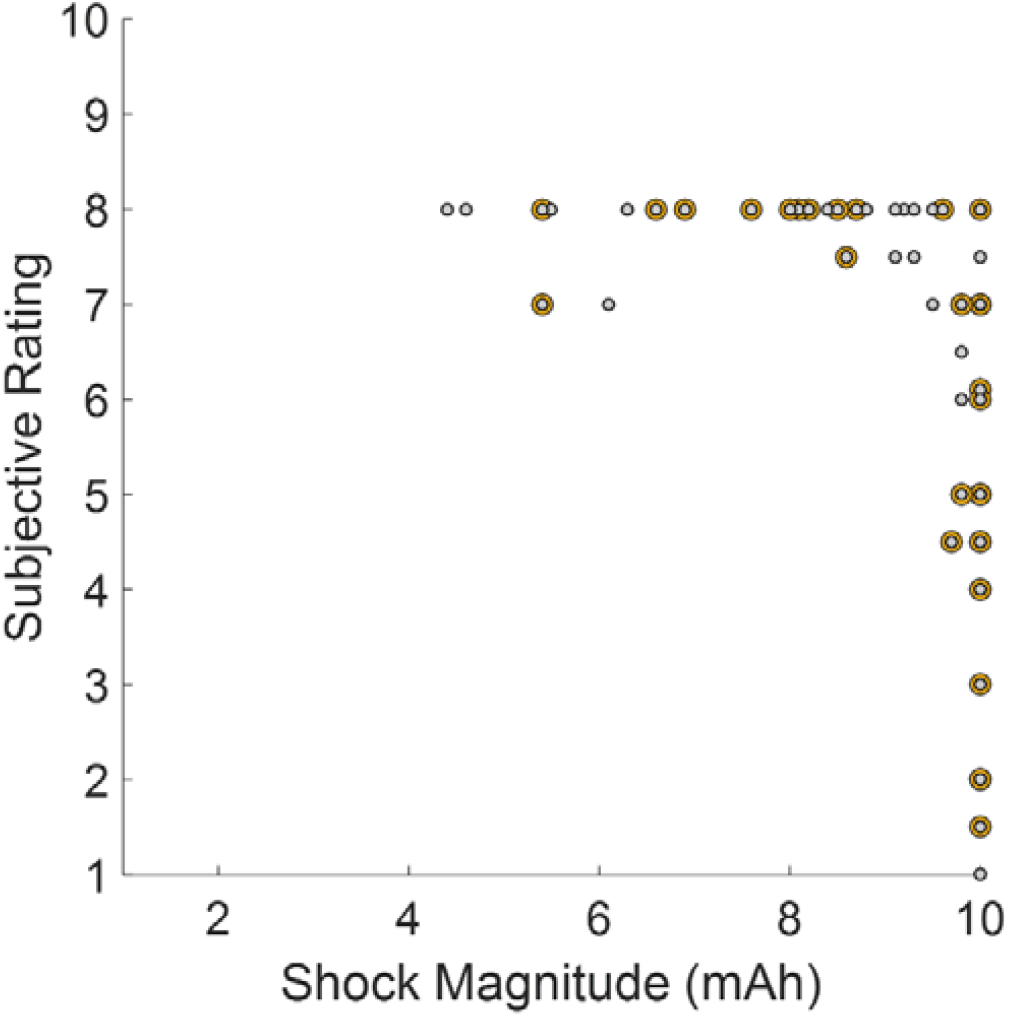
Graphical illustration of electric pain magnitudes and subjective pain ratings. Ketamine participants orange markers, placebo participants grey markers. As expected, there was a significant negative correlation between subjective pain ratings and electric shock magnitude (r(64)= -.495, p<.001) as it becomes unethical to increase shock magnitude when participant reports that the given electric pain magnitude in mAh meets the 80% tolerability threshold.

Intriguingly, right Hb activity in response to expectation of aversive outcomes (**Figure 3A**) also predicted subsequent preference behaviour for fractals specifically associated with loss outcomes (r(63) =0.246, p=0.046). By contrast, no relationship between right Hb activity and subsequent choice preference was observed for fractals associated with win outcomes (r(63) =.126, p=0.31).

## Discussion

In this study, we investigated the translation of findings from rodent models showing that direct ketamine infusion into the Hb suppresses burst firing associated depressive-like behaviours. Using a Pavlovian learning paradigm with peripheral electrical pain stimuli inside a 7-Tesla MRI scanner, we tested whether intravenous ketamine (administered 24 hours prior) modulates human Hb activity. We found that participants in the ketamine group exhibited overall reduced Hb responses during the expectation of aversive outcomes (**Figure 2B, 3A**). During observational learning of CS-outcome associations, the effects of ketamine were seen more generally in the aversive condition (across electric pain and monetary loss). Within the right Hb ROI, ketamine reduced Hb activation also for aversive versus positive outcomes compared to the placebo group (**Figure 4A**), indicating attenuated right Hb engagement after ketamine. These results provide preliminary evidence that ketamine modulates Hb function during aversive processing in humans, aligning with mechanistic predictions derived from preclinical studies. Together, these findings suggest that ketamine attenuates human Hb activity in a manner analogous to rodent models. Replication in clinical populations will be essential to establish the relevance of this mechanism for depressive pathology and treatment response.

This work translates a key finding from rodent models to humans for the first time. Preclinical studies converge on the lateral Hb as a key locus for ketamine’s rapid antidepressant-like actions. In rodent depression models, lateral Hb neurons show increased NMDAR-dependent burst firing relative to tonic or silent states^7,22^. Systemic ketamine blocks open NMDAR channels, preferentially silencing neurons with high basal burst rates, thereby reducing overall lateral Hb firing^10,23^. This state-dependent open-channel block persists beyond plasma clearance, as ketamine can remain trapped in the channel pore, yielding a sustained suppression of the lateral Hb bursting for approximately 24 hours after a single dose in chronically stressed mice, an effect that is not seen with saline and which is largely dissipated by 72 hours^10,24^. Behaviorally, these circuit changes track with rapid improvements in depression-like behavioural assays, including an association with reduced immobility on the forced swim test and increased sucrose preference, whether ketamine is given systemically (e.g., 10–25 mg/kg i.p.) or infused bilaterally into the lateral Hb^25-27^. On the other hand, previous human neuroimaging studies implicating the Hb as a centre for ketamine activity are suggestive but methodologically limited. One study^13^ exploring the activity of the Hb during aversive processing in MDD showed that depressed subjects have abnormal phasic Hb responses to cues that predict upcoming punishment (electric shocks). Further, in both MDD and healthy controls, a smaller Hb volume was associated with greater anhedonia^13^. One PET study reported reduced Hb glucose metabolism after a single sub-anesthetic ketamine infusion^28^. In this work, regional cerebral glucose metabolism was measured before and after a single ketamine infusion (0.5 mg/kg IV over 40 min) in 20 unmedicated patients with treatment-resistant MDD. Whole brain metabolism did not change, but regional decreases in metabolism were seen following ketamine, including in the right Hb, and the rapid antidepressant response was associated with reduced right Hb and prefrontal cortical metabolism. A separate 3T resting-state fMRI study reported that increases in right Hb functional connectivity with frontal, temporal, parahippocampal, and occipital regions correlated with symptom improvement following ketamine treatment^29^. However, these studies are constrained by insufficient spatial resolution for precise Hb localisation, increasing susceptibility to partial-volume contamination from adjacent thalamus and ventricular regions, and by the absence of placebo or non-clinical control groups, which restricts causal interpretation of ketamine’s Hb effects. Importantly, our study overcomes these limitations by leveraging high-field 7 T, task-based fMRI, a priori Hb ROIs, and a randomized, placebo-controlled design. In addition, the paradigm we used explicitly probes aversive processing (via monetary loss and electrical pain) interleaved with reward, enabling valence-specific contrasts and evaluation of cross-valence coupling. Collectively, these methodological features provide the most direct, high-resolution human test to date of ketamine’s predicted attenuation of Hb responses to negative outcomes.

Further, the Hb is increasingly recognized as a key structure in the neurobiology of MDD, owing to its central role in regulating negatively motivated behaviour^11^ and its strong inhibitory control over midbrain monoaminergic systems^30,31^. The Hb is anatomically and functionally connected to both the ventral tegmental area (VTA) and the raphe nuclei, enabling it to suppress dopaminergic firing and influence affective states such as anhedonia and low motivation^11,32-35^. It responds not only to primary aversive events but also to cues predicting negative outcomes and to omission of rewards, making it a critical hub for processing “anti-reward” signals^11,12,36^. Preclinical models implicate Hb hyperactivity in depression-like behaviours^27^, yet human imaging data are mixed. In particular, a 3 T fMRI Pavlovian conditioning study^13^ showed reduced Hb activation to shock-predicting cues in MDD (n=25) compared with healthy controls (n=25), opposite to the hyperactivity typically reported in rodent models. This inconsistency highlights the need to clarify how the Hb contributes to depressive pathophysiology and whether treatments such as ketamine normalize Hb function or act via broader Hb-centred circuits. Addressing these questions could refine mechanistic models of MDD and inform development of novel, circuit-targeted antidepressant strategies.

Finally, while our work is consistent with a previous preclinically established mechanism by which ketamine acts on lateral Hb neurons, it is important to highlight several differences. Preclinical studies have primarily used congenitally learned helplessness rats^7^ exhibiting depressive-like behaviours at the time of ketamine infusion, whereas our sample comprised healthy volunteers without clinically relevant past or current mood symptoms. In rodents, ketamine’s effects emerge specifically when lateral Hb neurons display a burst firing pattern of activity, an activity state that occurs only following a stress or negative mood induction associated with depressive-like behaviours. In contrast, our event-related fMRI paradigm is more similar to optogenetic manipulation of this circuitry^37,38^ as it was fine-tuned to heighten Hb activity at specific time points, for example when participants received pain stimuli versus when they won money in the task (**Figure 4A**). From a translational neuroscience perspective, the convergence between these findings in humans and preclinical models, despite methodological differences, provides preliminary support for a central role of Hb-related mechanisms in ketamine’s actions. Future work should examine these circuits in clinical populations to strengthen translational relevance.

Our results also highlight a potential cognitive pathway by which ketamine may act as a rapid antidepressant. We used a Pavlovian aversive conditioning paradigm shown to be a reliable probe of the Hb circuitry in humans^13,14^; however, this approach lacks behavioural choice during learning, and successful learning must instead be inferred from participants’ subsequent choices in a preference test. This preference test was administered approximately half an hour after the corresponding learning blocks, a delay that likely increases reliance on hippocampal long-term memory systems^39,40^. We observed that participants who had reduced habenula activity, more common in the ketamine group (**Figure 3A**), were more likely to prefer fractals associated with pain (**Figure 3B**). This relationship, based on Hb activity during the expectation phase, may reflect participants maintaining the expected outcome probabilities in their working memory, that is, the same information which is demanded while making value-based decisions^41^ during the preference test. Although exploratory, this suggests that ketamine may reduce the behavioural impact of negative affective experiences. These findings may inform reverse-translational work, as ketamine’s effect on depressive behaviours and negative affective memory^42^ have not yet been examined together in rodents. Future studies using instrumental learning in dynamically changing environments^43,44^, such that the habenular aversive-learning circuitry would be continuously engaged during the task^8^, may allow more precise isolation of ketamine’s cognitive effects.

A key limitation is that the cohort comprised healthy volunteers. Although this design provides a symptom-free characterisation of ketamine’s effects on right Hb function, it restricts generalisability to MDD and prevents evaluation of whether habenular changes mediate antidepressant or anti-anhedonic outcomes. Future work should include high-resolution, task-based 7T fMRI in MDD to link Hb modulation to clinical improvement. Establishing this brain–behavior link can clarify the neural basis of rapid antidepressant response, identify predictors of durability, and inform optimization of long-term therapeutic strategies (e.g., dosing schedules, adjunctive interventions).

In conclusion, this study provides evidence that ketamine, administered 24 hours before testing, reduces right Hb responses to aversive stimuli in healthy participants. These findings represent the first human translation of preclinical findings and support the idea that ketamine’s rapid antidepressant effects may be mediated, at least in part, by modulation of Hb activity. Future research in clinical populations is needed to strengthen and extend these findings. A clearer understanding of ketamine’s mechanism of action as a rapid-acting antidepressant will refine pathophysiological models of MDD, guide target identification/validation for drug development, and inform mechanism-based treatment decisions.

## Materials and Methods

### Participants

Seventy participants between the ages of 18 and 45 years were enrolled in this study. All had normal or corrected to normal vision, had no present or past neurological or psychiatric diagnosis, and provided written informed consent to participate. Healthy mood history was confirmed by a Structured Clinical Interview for DSM-5 Disorders (Research Version; SCID-5-RV) which was conducted by EP, PAH and SC. EP and PAH interviewed participants in consultation with SC and PJC who are fully qualified consultant psychiatrists. Additionally, all participants underwent a physical examination, hematological, biochemistry and thyroid function laboratory testing, urine toxicology and pregnancy (as applicable) testing, and an electrocardiogram (ECG). Subjects with concomitant unstable medical illnesses or previous use of phenylcyclohexyl piperidine (PCP) or ketamine were excluded.

The study was approved by the University of Oxford Central University Research Ethics Committee (CUREC, R73654/RE001), and a signed informed consent document was obtained from all participants. The study was registered at clinicaltrials.gov (NCT04850911) and on the ISRCTN registry (ISRCTN13629652).

The reported Pavlovian experiment was a part of a larger trial that spanned across approximately 8 weeks from the time of clinical interview. Participants were compensated £250 for their time; they did not receive any additional reimbursement in relation to their performance in the current experiment. In total, data were not included for 4 participants: 1 due to excessive wraparound artefacts affecting the participant’s T1 structural images, 1 due to a wide scale power outage which affected Oxford at the time of scanning, and 2 due to scanner failure. The final sample comprised 66 participants (see Table 1 for demographic characteristics and baseline self-report questionnaires, including subjective state ratings).

**Table 1.** Demographic features of the study cohort and descriptive statistics relating to key mood symptoms such as depression (BDI: Beck Depression Inventory), state and trait anxiety (STAI) and anhedonia (TEPS anticipatory (Ant) and consummatory (Con) subscales).

|  | *Ketamine (n=35)* | *Placebo (n=31)* |
| --- | --- | --- |
| *Age, Mean (SD)* | 23.46 (5.3) | 23.52 (4.5) |
| *Gender, Male, n (%)* | 19 (54.3) | 15 (48.4) |
| *Race/Ethnicity, n (%)* |  |  |
| *White/Caucasian* | 29 (82.9) | 23 (74.2) |
| *Mixed/multiple ethnic groups* | - | 5 (16.1) |
| *Asian/Asian British* | 4 (11.4) | 3 (9.7) |
| *Prefer not to say* | 1 (2.9) |  |
| *Educational Attainment, years (SD)* | 16.89 (2.61) | 17.39 (2.629) |
| ***Questionnaire Measures [Mean (SD)]*** |  |  |
| *STAI-T* | 29.82 (4.979) | 29.32 (9.253) |
| *STAI-S* | 28.2 (6.365) | 26.97 (8.758) |
| *BDI* | 1.71 (2.037) | 2.06 (4.697) |
| *TEPS Ant* | 44.71 (5.165) | 46.29 (4.818) |
| *TEPS Con* | 40.74 (3.830) | 41.16 (4.810) |

## Experimental Procedures

### Ketamine infusion

Following screening, subjects who met all eligibility criteria returned to the Clinical Research Facility (CRF) to undergo randomisation. On arrival, participants were briefly reviewed to ensure the inclusion/exclusion criteria were still met and concomitant medication were checked by qualified medical personnel. After confirming that participants were nil per os (NPO), tested negative on urine pregnancy (as applicable) and toxicology tests, the subjects were randomized by an unblinded member of the study team. Participants were randomised to either ketamine hydrochloride (0.5 mg/kg) or placebo (saline NaCl 0.9%) in blocks of four according to a 1:1 randomisation schedule generated using the website sealedenvelope.com. Randomisation was stratified by gender (Male, Female). To ensure double-blind administration of study treatment, the randomisation codes, study drug, and placebo (saline) were stored at the Clinical Research facility (CRF) at the Warneford Hospital, Oxford, and packaging and labeling of test (ketamine) and control (saline) treatments were identical to maintain the blind.

The study drugs were administered according to well-established standard protocols. An indwelling catheter was placed in the antecubital vein, and the drug was administered over 40 min at a constant rate (60ml/hour) via an infusion pump. Pulse, blood pressure, and oxygen saturation were monitored every 10 minutes during infusion and every 15 min after the infusion for at least 1 hour. One hour after the cessation of the infusion, the participants completed another battery of questionnaire measures (see Table 1) and tasks, to be able to assess differences in domains such as self-reported side effects, current mood and anxiety symptoms. At the end of the visit, a doctor discharged the participant and completed Readiness to Leave checklist. Participants were advised not to drive, cycle or operate machinery until they have had a restful sleep or the following morning, whichever was the later. They were also advised not to drink alcohol or take any other medication, or not to sign legal documents on the day of the infusion visit. Participants received 24-hour contact information of the medically qualified personnel from the Biomedical Research Centre on-call rota who they could contact if any questions/concerns arose following the infusion.

### Pain calibration

Electric pain was delivered to the left hand (fascia over adductor pollicis muscle) via a custom magnetically shielded electrode, using a single 1,000-Hz electrical pulse. Subjects underwent a thresholding procedure to control for heterogeneity in skin resistance and pain tolerance. Prior to the scanning, shocks were administered sequentially with step increases in amplitude, and subjects provided verbal ratings of each shock on a scale from 0 (not painful) to 10 (terrible pain/pain equivalent of burning one’s hand severely/pain that would cause me to twitch and move inside the scanner) without seeing the shock magnitude. The level of shock delivered in the experiment was set to 80% of the maximum tolerated for each individual. The average shock magnitude was [mean±SD] 9.01±1.51mA. Ketamine group had marginally lower subjective pain ratings just prior to the beginning of the experiment (5.90 vs 6.81, t(64)=-1.877, p=.065, Figure 5). These subjective pain ratings were used as covariates while investigating the drug effects on neural activity. During the experiment, the electric shocks were delivered by using a Digitimer DS8R direct current stimulator which was triggered by custom MATLAB scripts.

### Pavlovian conditioning paradigm

We adapted a Pavlovian paradigm which was originally implemented by Lawson and colleagues^14^ (**Figure 1**). Abstract fractal images (i.e. the conditioned stimuli — CS) were probabilistically paired with win, loss, shock, or neutral outcomes. There were seven CSs per each block of the task, associated with the following fixed outcome associations: 80% chance of £1 win, 20% chance of £1 win, 80% chance of £1 loss, 20% chance of £1 loss, 80% chance of shock, 20% chance of shock, and 100% no outcome (neutral). All the stimuli were used in a subsequent preference test which was administered behaviourally. CSs were luminance-matched and assigned to conditions randomly across subjects. On trials where the reinforcing outcome (win, lose, or shock) was not presented, and on neutral trial outcomes, the word “nothing” was presented on-screen. The timeline of the experimental task is shown in Figure 1. On each trial, subjects initially saw a fixation cross, which remained on-screen for the entire trial; the CS appeared on average after 1000 ms, remaining on-screen until the end of the trial; and the outcome was presented on average 1500ms following the CS onset. CS/outcome sequences were pseudo randomised. Temporal jittering and decorrelation of task epoch durations were implemented to enhance the efficiency of the fMRI design.

Pavlovian/observational nature of the task meant that there was no behavioural input from the participants that can influence the outcomes. In order to make sure that participants paid adequate attention to the task and to minimise the possibility of participants falling asleep inside the scanner, on 20% of trials, the fixation cross present in the center of the screen turned from black to red for 300 ms during CS presentation (but before outcome), and participants were instructed to respond via a button press as quickly as possible. They were explicitly instructed that their responses made no difference to the outcomes they received. In total, 420 trials were presented over three blocks, which lasted 9.3 min each.

### Memory-guided preference task

In order to assess participants observational learning performance, a preference test was administered after the scanning session is completed. The preference test involved pairwise comparisons between all the shapes which were presented within a given block and the trials were pseudo-randomised such that shapes from the 1^st^ block of the scanning task were presented initially, and the shapes from the final block of the task were presented last in order to control for primacy/recency effects. Participants were asked to declare their preferences between CSs they learned about throughout the Pavlovian learning task, and they were explicitly informed that their performance in the preference test would not affect their final reimbursement. All the experimental tasks were administered using Psychtoolbox 3.0 running on MATLAB. Approximately, participants completed the preference test between the fractals they learned about after 30 minutes from the completion of the corresponding learning block.

### fMRI acquisition

MRI data were acquired with a Magnetom 7T scanner (Siemens Healthineers AG, Germany) fitted with a single channel transmit / 32 channel receive head coil (Nova Medical Inc., United States). T2*-weighted images were obtained using a 2D multiband EPI sequence^45^ with 1.2mm isotropic resolution and 192 x 192 x 60 mm^3^ field of view (50 slices, TR = 1375 ms, TE = 19 ms, flip angle = 60°, multiband factor = 2, PAT factor = 3, partial Fourier = 6/8, BW = 1420 Hz/px). The field of view was centred manually in line with the habenula in each subject. 418 volumes were collected in each block, and the first five were discarded to account for T1 saturation effects. To allow distortion correction of the functional images, a GRE field map was acquired with 2mm isotropic resolution, 192 mm x 192 mm field of view, 77 slices, TR = 620 ms, TEs of 4.08 ms and 5.1 ms, and a flip angle of 39°. To help with registration, a single EPI volume was acquired with increased whole brain coverage (120 slices, TR = 3298 ms) but otherwise identical parameters and identical shim settings as the functional images. Peripheral pulse and respiratory effort were recorded during the EPI scans using a photoplethysmograph and pneumatic chest belt connected to a modular data acquisition system (MP150, BIOPAC Systems Inc., USA) These were used to correct for pulse- and respiration-related artifacts during analysis through physiological noise modelling. High-resolution T1-weighted images for registration of functional images were acquired with an MPRAGE 3D sequence. This had 0.7 mm isotropic resolution, 224 mm x 224 mm x 179.2 mm field of view, TR = 2200 ms, TE = 1.65 ms, TI = 1050 ms, flip angle = 7°, and PAT factor = 2. T1 structural images contained 256 slices.

### fMRI analysis

MRI data were analysed using FSL (FMRIB Software Library v6.0; https://fsl.fmrib.ox.ac.uk/fsl). Structural anatomical scans were brain-extracted and B1-bias-field corrected using the *fsl_anat pipeline*. fMRI data were preprocessed and analysed using FEAT (FMRI Expert Analysis Tool, version 6.0, part of FSL). Preprocessing steps included: 1. brain extraction using BET to remove non-brain tissue; 2. motion correction using MCFLIRT (6 parameters, Jenkinson et al., 2002), along with identification and exclusion of outlier volumes exhibiting excessive motion using *fsl_motion_outliers*; 3. spatial smoothing using a 3-mm full-width at half maximum (FWHM) Gaussian kernel to enhance signal-to-noise ratio while preserving spatial specificity in small structures such as the Hb; 4. unwarping using field maps to correct for geometric distortions in the echo-planar images; 5. high-pass temporal filtering with a 60-second cut-off to remove low-frequency drifts. Preprocessing of all fMRI data was conducted throughout the trial by study team members who were blinded to treatment allocation. Functional images were registered linearly to each participant’s structural image via an intermediate high-contrast whole-brain EPI image using boundary-based registration (BBR), optimizing anatomical alignment for regions of interest.

### Model-based analysis

*FSL’s FEAT was used to analyse all 1*^*st*^ *(individual runs), 2*^*nd*^ *(summary within-subjects) and 3*^*rd*^ *(between-group) functional data*. In the 1^st^ GLM that we fitted to the data, our main regressor of interest was the expected value of CSs timestamped to the presentation of abstract fractals. Below we describe how the expected value regressor was generated.

Given uncertainty about whether ketamine would influence instrumental learning, instead of using a Rescorla-Wagner model^46^ with an assigned learning rate value [which may in fact be different between ketamine and placebo groups], we decided to use a Beta distribution update rule that can track probabilities of outcomes optimally. Let ***O*** be the outcomes observed during the task, ***O***∈(win, loss, shock, null), and let estimated probability of an outcome be 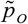. In a Beta distribution,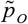 is updated via under the control of two parameters, namely **α**and **β**:

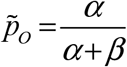

where on each trial α parameter for a given fractal type is updated as α+1 if an outcome is observed, or β parameter is updated as β+1 if the probabilistic outcome returns null. On trial 1, the update rule starts with values α=1 and β=1, meaning 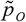 on trial 1 would be 0.5. This approach would generate identical learning trajectories for each participant, gradually reaching an asymptote towards the true probability of an outcome (e.g. starting from 0.5 and gradually approaching to 0.8 for high probability fractals). In the current experiment, for the number of trials for each fractal, Rescorla-Wagner equivalent of these learning trajectories would have an approximate learning rate of 0.124.

In the 1^st^ event-related neuroimaging model we aimed at improving the precision of these regressors [which are otherwise identical for each participant] by further convolving them with the single-subject decision outputs from the preference test that we used as a proxy for perceived valence of outcomes (e.g. if the shock fractals were preferred less relative to loss fractals, it would also mean that they should be perceived more negatively). This allowed the expected probability regressors to be linearly scaled from most appetitive to most aversive within a single regressor; while also reflecting between-subject variability in how the outcomes were perceived (e.g. for some participants fractals associated with 80% loss outcome were preferred less relative to fractals associated with 20% shock outcomes, indicating certain indifference points in choice preference). Therefore, this model tracks regions correlating positively or negatively with salient expectations.

A 2^nd^ event-related neuroimaging model which was constructed in relation to our preregistered and valence-specific outcome measures particularly related to fractals associated with electric pain. The estimated probability of outcomes for each outcome type were entered as distinct, valence-specific regressors (i.e. 4 expectation regressors for each of the CS types: win, loss, shock and neutral). Valence-specific outcome regressors were also entered into the model, and these were parametrically modulated by participants’ preference for each type of outcome (i.e. 4 outcome regressors for each of the US types: win, loss, shock and neutral).

Where applicable, all 1^st^ level models accounting for single subject neural activity in individual task runs were subjected to a 2^nd^ level analysis in FSL’s FEAT to get the average neural activity across all task blocks, and a further 3^rd^ level analysis directly comparing neural activity differences between placebo and ketamine groups.

### Model free analysis

In the model-free analysis, first-level analyses were conducted separately for each of the three functional runs using FEAT (FSL v6.0). First-level event-related GLM also included regressors generated by FSL’s physiological noise modelling GUI, to improve image quality and signal-to-noise ratio. Regressors were included for both CS and outcome events to model neural responses associated with anticipation and outcome processing. CS-related activity was modelled using four regressors per run (win, loss, shock, and neutral), along with additional contrasts reflecting grouped valence conditions: negative (shock + loss), positive (win) vs. negative (shock + loss), negative vs. positive. The first trial in which participants observed each CS fractal for the first time was excluded from the analysis to control for novelty-related responses. Outcome-related activity was modelled using regressors for each outcome type (win, loss, shock, and neutral) and similar valence-based contrasts (negative, positive vs. negative, negative vs. positive). All regressors were convolved with a canonical double-gamma hemodynamic response function (HRF), and temporal derivatives were included.

For each subject, contrast estimates from the three runs were combined using a fixed-effects model to produce an average contrast per condition using a second-level FEAT analysis (i.e., 2^nd^ level analysis).

### Statistical analysis

Higher (i.e. 3^rd^) level group analysis was carried out using FSL’s tool for nonparametric permutation inference *Randomise* (5000 permutations)^47^ to assess general effects of task-relevant contrasts on both groups, as well as test for group differences. Statistics were assessed using the threshold-free cluster enhancement method with family-wise error correction of 0.05 (or 0.95 threshold within randomise)^48^. Small-volume correction (SVC) was applied using a right habenula (Hb) mask derived from the high-resolution probabilistic subcortical atlas of Pauli, Nili, & Tyszka (2018). Specifically, we extracted the right Hb label from their atlas (MNI152 space), thresholded the probability map at ≥50%, and binarized it to create the ROI. All Hb inferences report ROI-level FWE-corrected p-values based on N = 5000 permutations. The GLM included 2 groups: placebo and ketamine. Contrasts were defined as placebo greater than ketamine, ketamine greater than placebo, and the mean across both groups to establish main effect of task. Significant brain areas were extracted for visualization using the *fslmaths* and cluster tools, with a threshold of 0.95 (based on the 1-p thresholding from randomise, described above).

All behavioral and follow-up neuroimaging analyses on the regression coefficients extracted from our ROIs were done in MATLAB. Repeated measures ANOVA (rmANOVA) models and partial correlations were fitted in SPSS version 29.0 (IBM). All post-hoc pairwise comparisons relate to appropriate two-tailed t-tests.

## Acknowledgments

We wish to acknowledge the support of the NIHR Oxford Cognitive Health Clinical Research Facility, the Oxford Health Biomedical Research Centre (BRC), and the Oxford Centre for Integrative Neuroimaging (OxIN, previously Wellcome Centre WIN) for their assistance in conducting this study.

## Supplementary Figures, Results and Discussion

**Table S.1.**
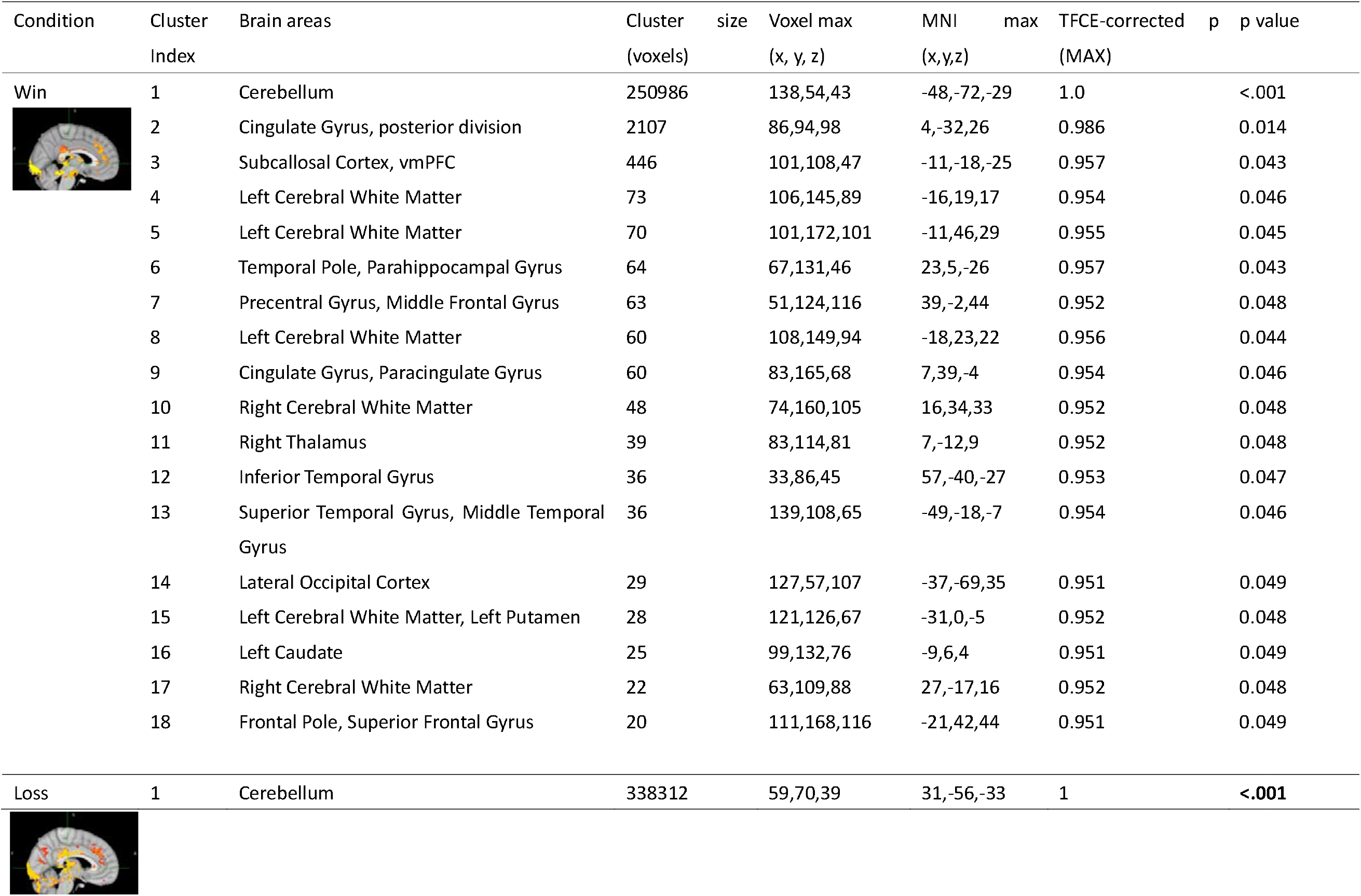
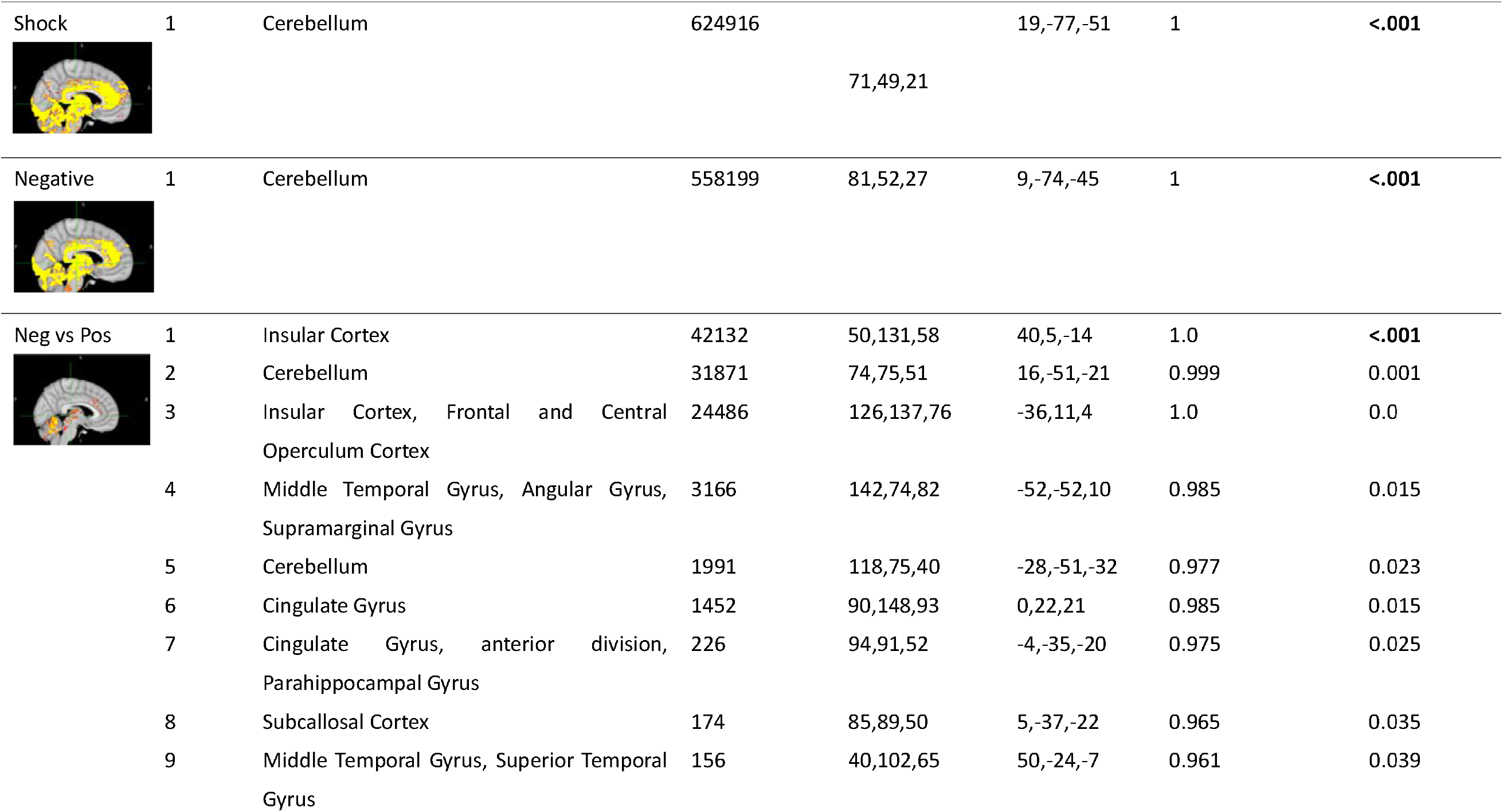
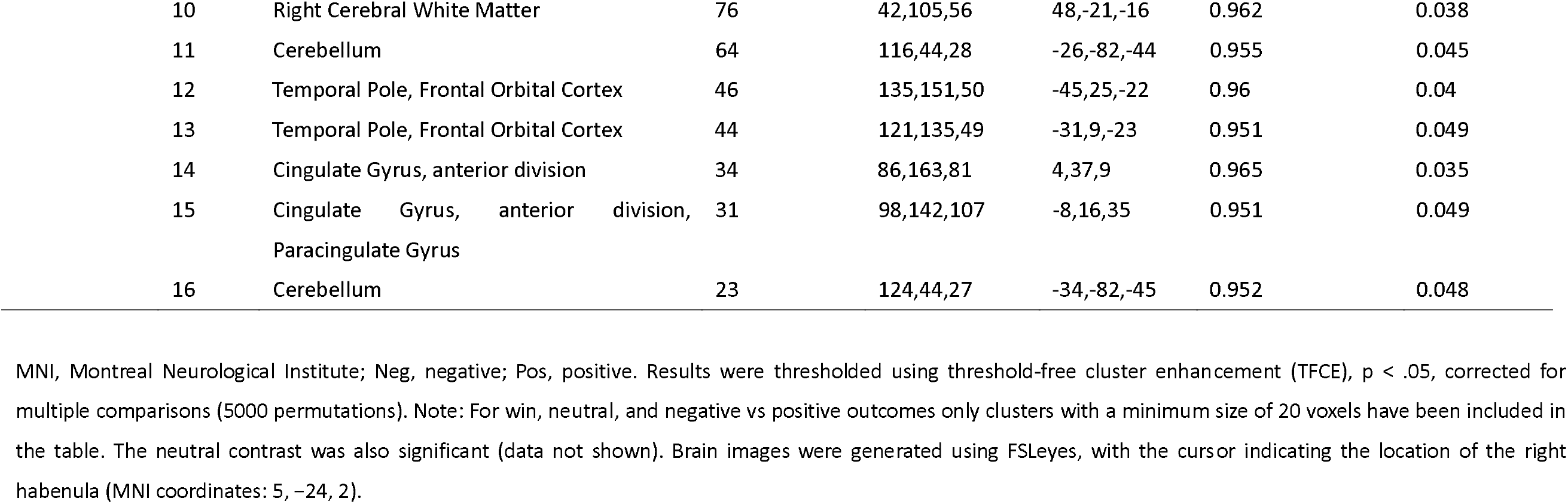
Cluster Peaks for Activations Related to the Unconditioned Stimuli (USs) in whole brain (WB) MNI, Montreal Neurological Institute; Neg, negative; Pos, positive. Results were thresholded using threshold-free cluster enhancement (TFCE), p < .05, corrected for multiple comparisons (5000 permutations). Note: For win, neutral, and negative vs positive outcomes only clusters with a minimum size of 20 voxels have been included in the table. The neutral contrast was also significant (data not shown). Brain images were generated using FSLeyes, with the cursor indicating the location of the right habenula (MNI coordinates: 5, −24, 2).

**Table S.2.**
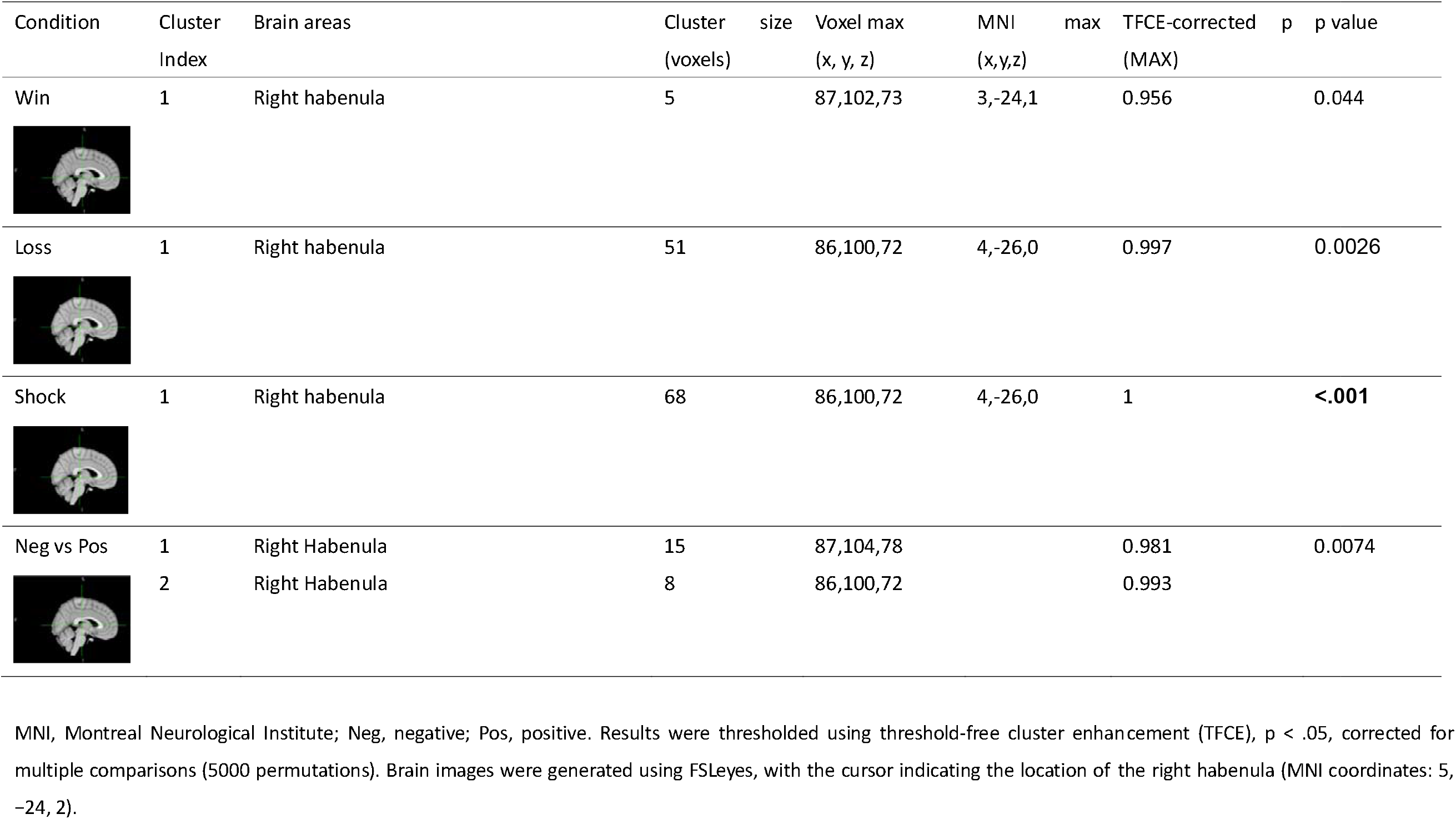
Cluster Peaks for Activations Related to the Unconditioned Stimuli (USs) in the Right Habenula (Hb) MNI, Montreal Neurological Institute; Neg, negative; Pos, positive. Results were thresholded using threshold-free cluster enhancement (TFCE), p < .05, corrected for multiple comparisons (5000 permutations). Brain images were generated using FSLeyes, with the cursor indicating the location of the right habenula (MNI coordinates: 5, −24, 2).

| Condition | Cluster Index | Brain areas | Cluster size (voxels) | Voxel max (x, y, z) | MNI (x,y,z) | max TFCE-corrected (MAX) | p | p value |
| --- | --- | --- | --- | --- | --- | --- | --- | --- |
| Win | 1 | Right habenula | 5 | 87,102,73 | 3,-24,1 | 0.956 |  | 0.044 |
| Loss | 1 | Right habenula | 51 | 86,100,72 | 4,-26,0 | 0.997 |  | 0.0026 |
| Shock | 1 | Right habenula | 68 | 86,100,72 | 4,-26,0 | 1 |  | **<.001** |
| Neg vs Pos | 1 | Right Habenula | 15 | 87,104,78 |  | 0.981 |  | 0.0074 |
|  | 2 | Right Habenula | 8 | 86,100,72 |  | 0.993 |  |  |

**Supplementary Figure 1.**
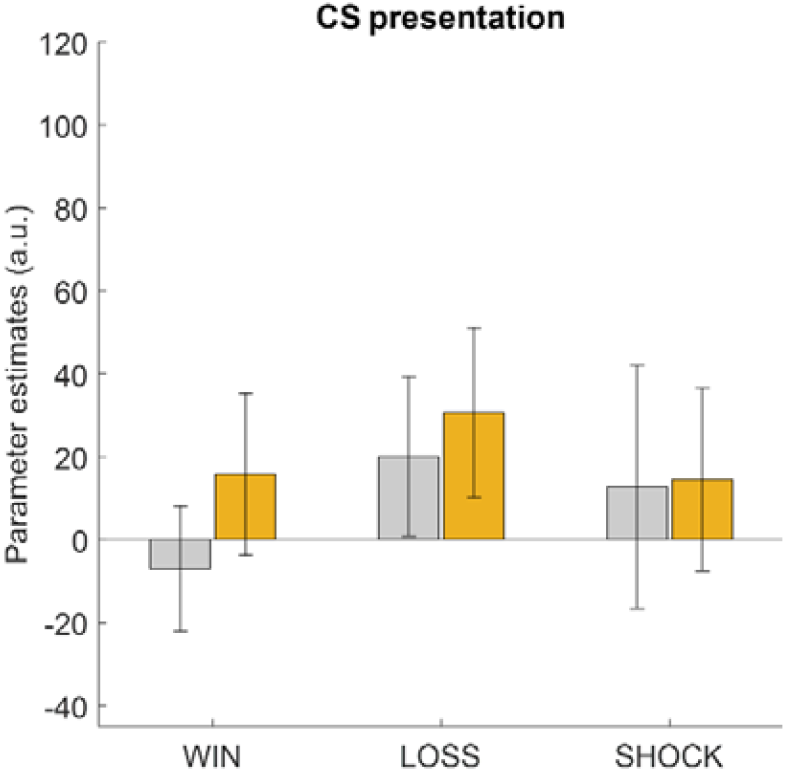
Decomposing the model-based MRI results of the expectation phase (from right Hb in Figure 2B & Figure 3A) down to its valence-specific components in a model-free manner fail to identify between group differences, indicating that ketamine acts on aversive outcome expectations generally, rather than expectations of loss or shock outcomes specifically. Error bars designate ±1SEM.

**Supplementary Figure 2.**
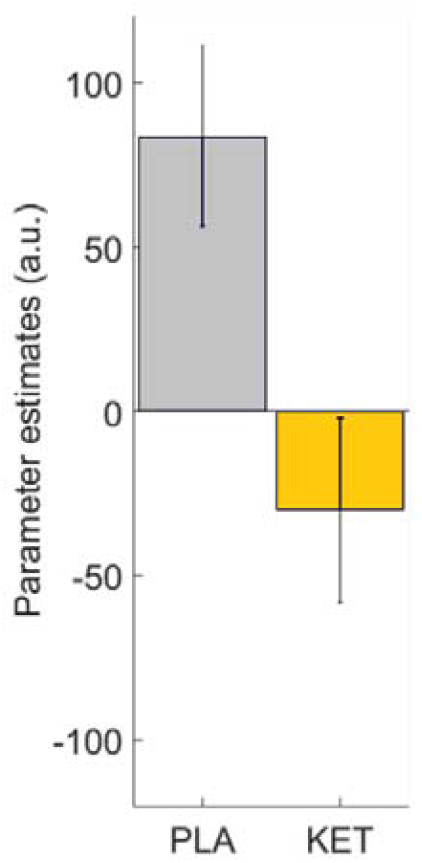
For the direct comparison of receipt of monetary win versus loss outcomes, the ketamine group (KET, in orange) had significantly higher activity in the right Hb relative to the placebo group (PLA, in grey). Error bars designate ±1SEM.

**Supplementary Figure 3.**
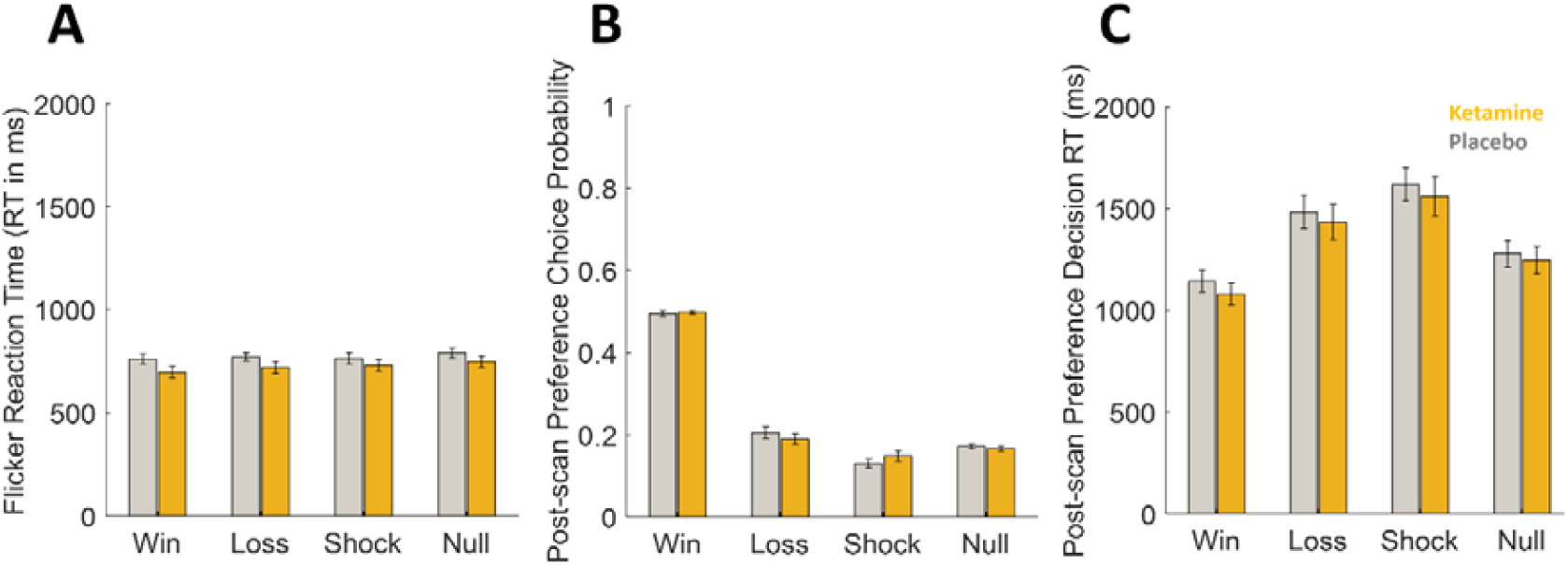
Summary of behavioural results. Globally, the groups were comparable in terms of reaction times (RTs) to the flicker detection element of the Pavlovian learning task and RTs during the preference test. The subtle differences observed in participant choice behaviour during the preference test were not significant (panel B). Error bars designate ±1SEM.

### The relationship between ketamine’s acute dissociative effects and right Hb response to expectations of aversive outcomes

It has been suggested that there may be a relationship between ketamine’s acute dissociative effects and clinical response^49^. Although our study was not conceptualised and/or powered to specifically test such associations, an analysis exploring this connection suggested a marginally negative relationship in the ketamine group between delta CADDS scores (Table 1, i.e. the difference between CADDS scores pre-and post-infusion), and habenula activity in (r(33)=-.28, p=.106).

### Ventral tegmental area encodes a meta learning signal about aversive environments

We further explored z-maps reported in **Figure 2A** at a WB p=.005 uncorrected threshold^14^ which helped us to identify activity in a key dopaminergic region that is in the vicinity of substantia nigra and ventral tegmental area (vTA) border (**Supplementary 3A**). This area is particularly important as previous monkey and rodent neurophysiology work suggested that habenula neurons work in tandem with neurons in the vTA^11,50,51^ during reinforcement learning. We extracted the regression coefficients associated with expected values of appetitive outcomes from this region using FSL’s *featquerry* and a 3mm spherical ROI drawn around the peak coordinates of the vTA activity. Subsequent analysis indicated a main effect of ketamine (F(1,64)= 5.009, p= .029, **Supplementary Figure 3B**), whereby ketamine augmented the activity of the vTA during expectation of positive outcomes. Gradually dampening activity observed across 3 runs of the task in the placebo group (**Supplementary Figure 3B**, grey bars) seems to suggest that through its temporal dynamics vTA may be encoding a meta-learning signal that guides a higher-level realisation that learning environments were globally aversive (i.e. two different aversive outcomes (monetary loss and electric pain) versus one type of positive outcome (monetary win), and one neutral outcome). One type of behaviour analysis that can support this interpretation is the relationship between reaction times (RTs) to the flickers which were presented during the task (**Supplementary Figure 2A**) and the activity in the VTA. Normally one would expect reaction times to improve over time, by general principles of participants learning what they need to do while performing the task. However, if the VTA activity is indicative of a meta-learning signal with which participants come to understand that the learning environments are overall more aversive, the relationship between RTs and the VTA activity should also change gradually from positive to negative, as responding to flickers in more aversive environments would be similar to a no-go conflict. In support of this interpretation, a similar pattern of results also emerged from a correlation analysis between the RTs with regards to approach to cues and the expected value regressor that underlies the VTA activity, computed specifically on flicker trials (**Supplementary Figure 3C**). Here, correlation coefficients gradually changing from positive to negative across runs indicate a higher level of response slowing which is to be expected in environments that are perceived to be aversive.

**Supplementary Figure 4.**
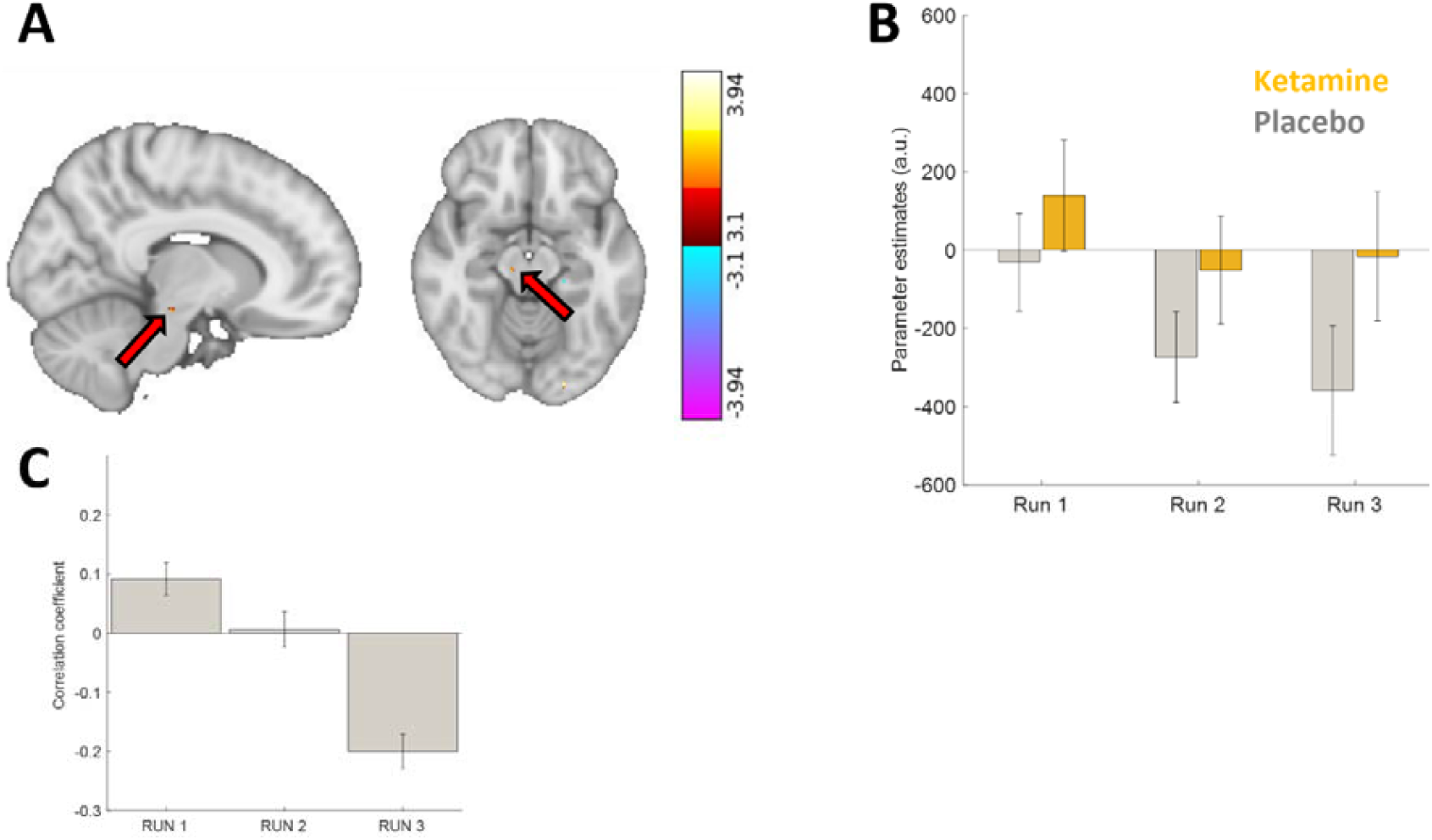
(A) Neuronal activity in the vicinity of ventral tegmental area and substantia nigra border shown across sagittal, coronal and axial views (highlighted by the red arrow, peak MNI: [11 -21 -14], z= 3.44. (B) Regression coefficients extracted from this region indicates a main effect of ketamine across all runs on neuronal activity, although the groups were not statistically significantly different within individual runs. (C) Correlation coefficients between VTA activity and reaction times to the flicker computed separately for each run. This analysis is only conducted for the placebo group (i.e. grey bars, also throughout the paper) to derive behavioural support complementing gradually dampening VTA activity also observed in this group. Linearly decreasing correlation coefficients from positive to negative across runs reflects the pattern of activity observed in the VTA. Error bars designate ±1SEM.

